# Internal perturbation reveals the flexible and adaptive nature of the coordination between decisions and movements

**DOI:** 10.64898/2026.07.23.740304

**Authors:** David Thura, Gislène Gardechaux, Clara Saleri

**Author notes:** Corresponding author:* David Thura Lyon Neuroscience Research Center – Impact team Inserm U1028 – CNRS UMR5292 – Lyon 1 University 16 avenue du Doyen Jean Lépine, 69675 Bron, France.

## Abstract

Although growing evidence indicates that decision-making and movement control are tightly coordinated, the functional significance of this coordination remains poorly understood. In particular, it is unclear whether changes in the animal’s arousal are sufficient to reorganize decision-action coordination in a manner that preserves behavioral efficiency. To address this question, we pharmacologically slowed behavior in rhesus monkeys performing a reaching-based foraging task while leaving task demands unchanged. Under this internal perturbation, prolonged decisions were initially associated with shorter movement durations, revealing a compensatory coordination that reduced the additional temporal cost of slower deliberation. As animals progressively adapted across sessions, decision durations decreased, reward rates increased, and compensatory coordination became less prevalent, giving way to patterns of decision-movement co-regulation. This observation suggests that compensatory adjustments are recruited transiently during adaptation rather than representing a constitutive mode of behavioral control. These findings provide causal evidence that an internally-induced slowing of decision-making is sufficient to reorganize the temporal relationship between decisions and movements. More broadly, they support the view that decision-making and movement execution are regulated as complementary components of a unified adaptive control process that flexibly adjusts the temporal organization of behavior in response to changes in internal state, thereby contributing to the maintenance of efficient reward acquisition.

## Introduction

Adaptive behavior depends on the ability to make appropriate decisions and to execute the corresponding actions efficiently. Although decision-making and movement execution have traditionally been investigated as separate processes, everyday behavior requires that they operate as components of a single behavioral sequence (Cisek and Kalaska, 2010). The time devoted to deliberation and the time required to execute the ensuing movement jointly determine how rapidly rewards can be acquired (Haith et al., 2012; Choi et al., 2014; Shadmehr et al., 2016; Morel et al., 2017). Consequently, understanding how these two processes are coordinated is essential for identifying the mechanisms that optimize behavioral performance.

A growing body of evidence in both human and non-human animals indicates that decision duration and movement vigor are tightly coordinated in a manner that promotes efficient behavior (for a review, see Thura et al., 2025). Under typical conditions, relatively short deliberation times are typically followed by vigorous movements, whereas longer decisions are accompanied by slower, less vigorous movements (Carsten et al., 2023; Kita et al., 2023; Saleri et al., 2026). Theoretical frameworks based on reward rate maximization provide a compelling explanation for this “default” mode of coordination (Shadmehr and Ahmed, 2020). Because both deliberation and movement contribute to the total duration of a behavioral phase, modifications in either component directly influence the long-term rate of reward acquisition. Consequently, an adaptive behavioral strategy should regulate decision and movement durations together rather than independently. According to this view, movement vigor is not determined solely by biomechanical or energetic considerations but also reflects the temporal costs incurred during decision formation. Likewise, decision policies should account for the anticipated costs of subsequent actions. The resulting behavioral coordination enables organisms to optimize overall performance in environments where time constitutes a limited resource.

Experimental studies have provided further support for this normative framework by demonstrating that the relationship between decision duration and movement vigor is flexible rather than fixed. When one component of the behavioral sequence becomes abnormally prolonged, compensatory adjustments may occur in the other component (Thura et al., 2014; Reynaud et al., 2020; Thura, 2020; Saleri Lunazzi et al., 2021; Saleri and Thura, 2024; Leroy et al., 2025; Saleri et al., 2026). For example, unusually long decisions can be followed by faster and more vigorous movements, thereby reducing the additional temporal cost associated with prolonged deliberation (Thura et al., 2014; Thura, 2020). Such compensation limits the increase in total trial duration and partially preserves the average reward rate (Saleri Lunazzi et al., 2021). These observations indicate that the sensorimotor system does not merely express stereotyped couplings between decisions and actions but dynamically regulates their coordination according to the temporal demands imposed by the task.

Although these findings strongly support the existence of integrated decision-movement control, an important question remains unresolved. The previous demonstrations of compensatory coordination have relied on manipulations that modified movement constraints, task demands, or decision difficulty. It is thus unknown whether an internal, experimentally-induced increase in behavior duration alone is sufficient to trigger adaptive modifications in the coordination between decisions and movements. In other words, if the decision becomes intrinsically slower, independently of task parameters, does the system spontaneously reorganize movement execution to adjust the coordination between the two processes so as to preserve the reward rate? Establishing this causal relationship is essential for determining whether the coordination between decision-making and movement control truly serves to compensate for temporal perturbations and maintain efficient behavior.

In the present study, we addressed this question by administrating a mild sedative to two out of three monkeys trained to perform a reaching-based patch foraging task. We asked whether a sedative-related increase in behavior duration alters the natural coordination between decision-making and movement in a manner consistent with the objective of optimizing reward rate.

## Material and Methods

### Animals, ethical framework, and testing context

Experiments were conducted with three rhesus macaque monkeys (Macaca *mulatta*): monkey G, female, ∼9 kg, 10-11 years old, right-handed; monkey B, male, 7-8 kg, 5-6 years old, left-handed; monkey D, male, ∼6 kg, 5 years old, left-handed. Data were collected over a total of 101 sessions for monkey G, 152 for monkey B and 36 for monkey D.

Ethical permission was provided by “Comité d’Éthique Lyonnais pour les Neurosciences Expérimentales” (CELYNE), C2EA #42, ref: C2EA42-11-11–0402-004, and by the French Ministry of Research and Higher Education (initially obtained in 2020, renewed in 2024). Monkey housing and care were in accordance with the European Community Council Directive (2010) and the Weatherall report, “The use of non-human primates in research”. Laboratory authorization was provided by the “Préfet de la Région Rhône-Alpes” and the “Directeur départemental de la protection des populations” under Approval Number E 69 029 0601 (renewed on May 28, 2026).

Behavioral data from monkey G and monkey B were collected head-fixed and simultaneously with electrophysiological recordings which required the manipulation of microelectrodes to isolate neuronal activity (which will be the topic of future publications). To this end, both animals were implanted with a titanium head fixation post and with one form-fitting recording chamber (Gray Matter Research). All surgical procedures were carried out under anesthesia and strict aseptic conditions. On the other hand, because monkey D was not yet implanted for neurophysiological recordings, his behavioral data were acquired under head-free conditions.

### Pharmacological procedure

All sessions involving monkey G (n=101) were conducted under light sedation, initially to ensure the animal’s well-being and reduce any stress associated with handling during electrophysiological recordings. To this end, Domitor^Ⓡ^ (medetomidine, 1 mg/ml) was administered prior to each session at a dose of 0.01mg/kg, by intramuscular injection. Medetomidine possesses sedative, analgesic, and muscle-relaxant properties. These properties result from the agonist effect of medetomidine on presynaptic and postsynaptic alpha-2-adrenergic receptors. Activation of these receptors leads to a reduction in the release and turnover of noradrenaline.

By contrast, most sessions performed by monkey B (n=133 out of 152) were made without sedation. To evaluate the impact of sedation on decision-action coordination and to compare the animals’ behavior with and without sedation, monkey B also performed 19 sessions under light sedation, with Domitor^Ⓡ^ administered prior to each of these 19 sessions at a dose of 0.00625mg/kg, by intramuscular injection. The first session executed under sedation occurred after 70 sessions performed without sedation. Then, during a typical week, monkey B underwent four sessions without sedation (Monday through Thursday) and one session with sedation (Friday).

In this sedative condition and for both animals, the behavioral task began when they were fully awake and attentive, typically about 10-30 minutes after the administration of Domitor^Ⓡ^ which is made in animals’ enclosure.

Finally, all monkey D’s behavioral data were acquired without the light sedation used for monkeys G and B.

### Experimental apparatus

The monkeys sat in a primate chair and were trained to perform a patch-foraging task based on planar arm movements. They held a lever in their dominant hand (right for monkey G, left for monkeys B and D) and a digitizing tablet (GTCO CalComp) continuously recorded its horizontal and vertical positions (100 Hz with 0.013 cm accuracy). The visual display was projected by a VIEWPixx monitor (VPixx Technologies; 120 Hz refresh rate) onto a mirror suspended 25 cm above and parallel to the digitizer plane (Figure 1A). Unconstrained eye movements and pupil area were recorded using an Eyelink 1000 (SR Research) infrared camera (data not shown).

**Figure 1-.**
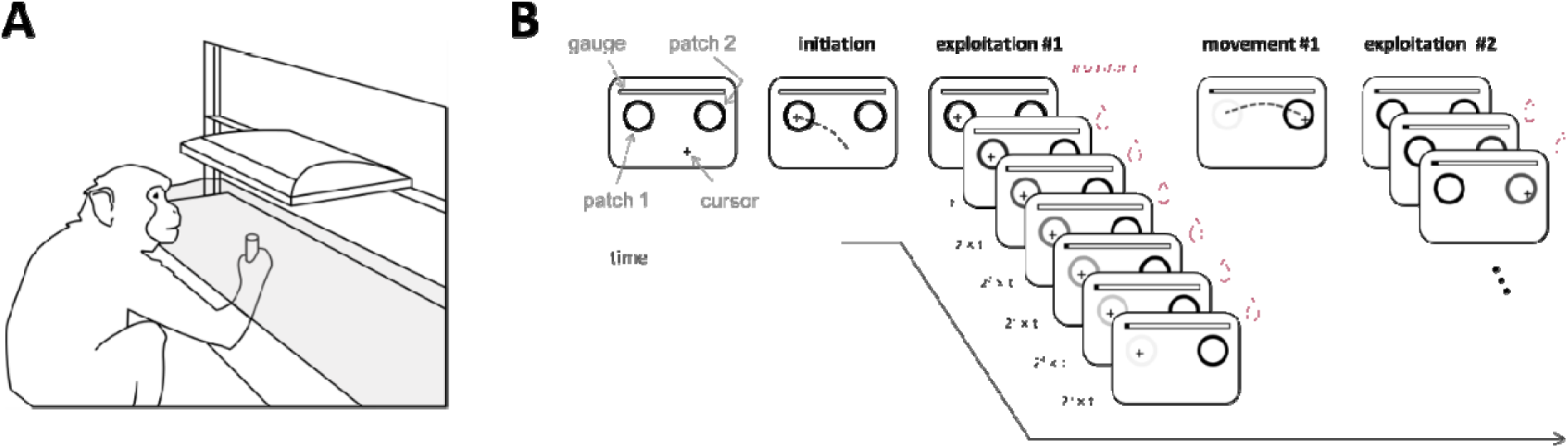
Experimental apparatus and behavioral task. A: Experimental apparatus. The monkeys sat in a primate chair and performed arm movements to control a lever held in their dominant hand. A digital tablet continuously recorded the horizontal and vertical positions of the lever. Visual stimuli and a cursor, which served as visual feedback of the lever, were projected onto a semi-transparent mirror positioned parallel above the digital tablet. B: Time course of a trial in the foraging task (see text for details).

### Behavioral task

The three macaques have been trained to execute a computerized, reaching-based patch foraging task (Figure 1B). In this task animals manipulated a lever and performed a reaching movement toward one of two possible visual targets (the reward sources, or “patches”) to collect drops of fruit juice. Patches were indicated by two black circles placed horizontally. The position of the lever was materialized on the screen by a black cross (hereafter referred to as the “cursor”). Once the lever was placed in a patch and held still for 100ms, juice drops were delivered at a rhythm that decreased over time, mimicking the exploitation of a depleting source of rewards. When the time interval between two drops was perceived to be too long, the monkeys could decide to leave the current patch by executing a reaching movement toward the other patch to start another exploitation. Reward rhythms and reaching movement costs were varied within and between blocks of trials, respectively (please see below and Saleri et al., 2026 for a description of the experimental conditions).

From the experimenter’s point of view, each drop of fruit juice collected by the monkey corresponds to a point. At the start of each session, the experimenter set a number of points that the monkeys had to collect from the two possible patches by alternating periods of static exploitations followed by arm movements executed to reach the other patch, in a back-and-forth manner (from the right patch to the left one, then from the left patch to the right one, and so on). On the top of the patches, a gauge that filled up as animals progressed in the task (in terms of accumulated rewards) was displayed. We assumed that efficient task performance involved animals adopting a strategy that maximized point accumulation while minimizing time spent in the experimental room.

At each exploitation phase, the monkeys could obtain up to 6 drops of juice, with the time interval between two being systematically multiplied by a factor of approximately two. This implies that the longer they stayed in a patch to exploit it, the longer they had to wait to get the next drop of juice. To visually reinforce the impression of exhaustion of a current source of reward, the color of the exploited circle gradually faded after each collected reward, disappearing completely by the 6th reward (Figure 1B).

In most patch foraging scenarios, the decision to leave a reward source to explore other opportunities depends on the current exploitation as well as the cost of the next exploration (Charnov, 1976; Hayden et al., 2011; Calhoun and Hayden, 2015; Barack, 2024). Therefore, by convention, a trial in this task was defined as an exploitation phase in one patch followed by an arm movement toward the other patch. This sequence of exploitation-movement was repeated by the monkeys as long as they were sufficiently motivated to perform it. If the gauge was full at the end of the session, they received, in addition to their liquid rewards accumulated during the session and their daily ration of accommodation, a “jackpot” reward, most often a whole fruit.

### Experimental conditions

In this foraging task, the exploitation of a patch is subject to a decreasing rhythm of rewards collected. As previously mentioned, the time interval between each of the 6 potential rewards increases, requiring the monkey to wait longer for the next one. The rhythm varied between 3 levels, from trial to trial. For the *Slow* rhythm, 17.2 ± 0.02 seconds (mean ± SD) were needed to obtain the 6 drops of juice, 11.4 ± 0.04 s for the *Medium* rhythm and only 5.7 ± 0.02 s for the *Fast* rhythm. Importantly, the rhythm of the next patch was unknown by the monkeys until they started to exploit it.

Moreover, the size and the distance of the patches were varied in three blocks of trials. In the *Accurate* condition, the patches were the smallest (1.25cm radius) and placed at a distance of 7cm from each other, encouraging slow and precise movements compared to a *Reference* condition in which the patches size was fixed at 2.25cm radius (but still located 7cm apart). By contrast, in the *Speed* condition, the size and distance of the patches (2.75cm radius and 10cm apart) encouraged the monkey to make more vigorous movements compared to the reference condition. During each session, monkeys performed blocks of approximately 50 trials per motor condition, continuing until achieving the daily objective of points.

The impact of reward rhythms and motor conditions on animal behavior lies beyond the scope of the present study and will therefore be described in future publications.

### Data analysis

A trial was considered valid for specific analyses if at least one reward was collected in the exploited patch, and if behavioral parameters (i.e. decision and movement durations) fell within ± 3 standard deviations of the session means, to exclude aberrant movements and exploitations. Among the excluded trials, almost all are those not initiated by the monkey due to a lack of stabilization in the patch or those for which the monkey left the patch before the arrival of the first drop of juice (between 10% and 15% of the trials, depending on the monkey).

The horizontal and vertical lever position data were used to calculate the different variables of interest described below. The kinematics data were first filtered using a 15th-degree polynomial filter and then differentiated to obtain a velocity profile. The onset and offset of each movement were determined by applying a velocity threshold of 2.5cm/s.

The exploitation phase (i.e. when the monkeys hold the lever inside one of the two patches and receive drops of juice at a decreasing rate) ends with the animal’s decision to leave the current patch and move to the other one. Thus, decision duration (DD) was calculated as the time between the stabilization of the cursor inside the current patch and the onset of the following movement executed to reach the other patch. For the “exploration” phase of each trial (i.e., the reaching movement), the duration of the movement (MD) was computed as the delay between movement onset and offset. The effectiveness of rewards collection, i.e. the reward rate, was quantified both at a local level, for every trial, and at the session level. The local reward rate (lRR), defined as the number of rewards collected per second of foraging, was computed for each trial by dividing the number of juice drops collected in a given trial by the sum of DD and MD. The average lRR for the session was then calculated by averaging these rates across all trials in the session. The global reward rate (gRR), defined as the number of rewards collected per minute of foraging, was calculated by dividing the total number of points accumulated over the session by the total duration of this session. Unlike the lRR, the gRR takes into account, in particular, potential pauses taken by the monkeys during the various experimental sessions.

### Statistics

We used MATLAB (MathWorks^®^) for all statistical analyses and the significance level of the tests was set at 0.05. Pearson correlations were computed to assess the relationship between DD and MD at the single-trial level (i.e. the degree of coordination between DD and MD). Pearson correlations were also used to test the evolution of decision and movement durations as well as reward rates (lRR and gRR) across sessions.

## Results

### General observations

Monkey G collected 1064 ± 154 drops of juice (mean ± SD) during the 101 sessions she completed, which required 235 ± 42 trials per session, with sessions lasting 29 minutes and 29 seconds on average. Monkey B collected 891 ± 195 rewards per session during the 133 sessions he performed without sedation, which required completing 297 ± 69 trials per session, with sessions lasting 17 minutes 3 seconds on average. During the 19 sessions he performed under sedation, monkey B collected 1367 ± 238 rewards per session, which required completing 390 ± 87 trials over 29 minutes and 17 seconds per session on average. Finally, monkey D collected 1316 ± 417 rewards through 405 ± 148 trials in 36 sessions, with sessions lasting 38 minutes and 40 seconds on average.

We first examined overall decision (DD) and movement (MD) durations in all three subjects, by separating the sessions conducted under sedation and those conducted without sedation for monkey B (Figure 2). As a reminder, monkey G performed the task only under sedation, whereas monkey D performed it only without sedation.

**Figure 2-.**
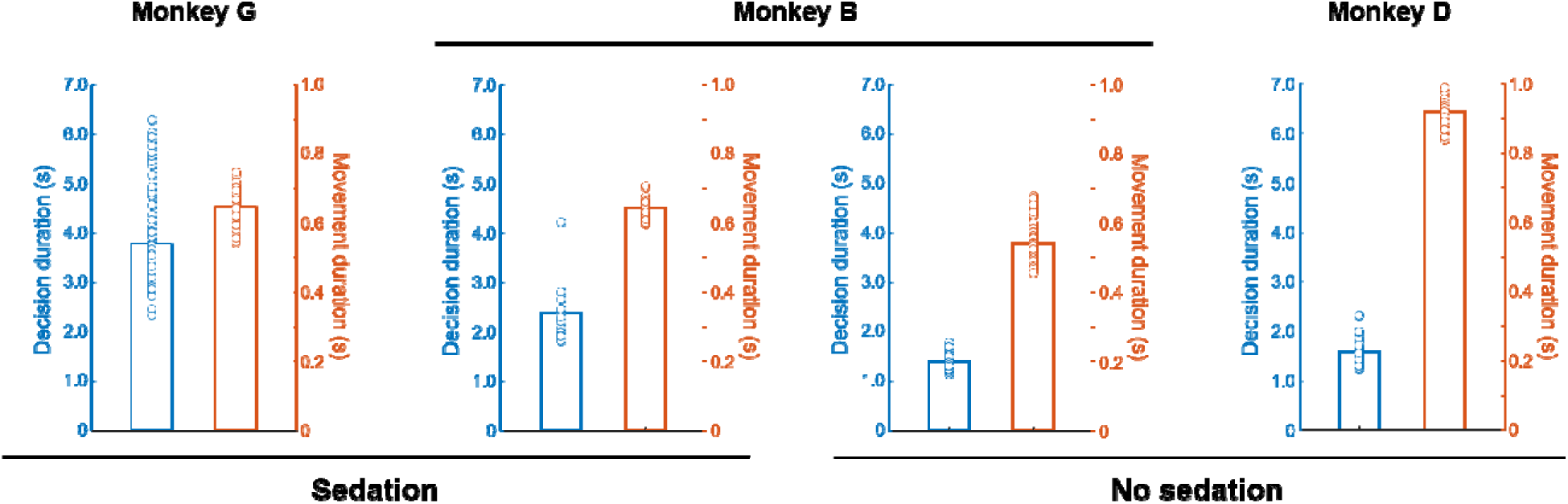
General behavior across sessions. Left panel: Decision duration (blue) and movement duration (orange) for monkey G who performed the task under sedation. Bar illustrate mean values across sessions and dots illustrate individual values for each session. Second column panel: Same as left for sessions performed by monkey B under sedation. Third column panel: Same as left for sessions performed by monkey B without sedation. Right panel: Same as left for sessions performed by monkey D without sedation.

Under sedation, monkey G exhibited the longest DD (mean ± SD: 3.79 ± 0.91s) compared to monkey B (2.37 ± 0.53s). Without sedation, the duration of decisions of monkey B decreased (1.40 ± 0.1s) compared to the sedation condition (2.37 ± 0.53s) and were shorter than those made by monkey D (1.57 ± 0.23s). Under sedation, monkey G and monkey B performed movements of comparable duration (647 ± 43ms and 641 ± 33ms, respectively). Without sedation, Monkey B showed the shortest movements (541 ± 49ms), followed by monkey D, who performed particularly long movements on average (916 ± 0.38s).

In sessions conducted under sedation with monkey G, a decrease in decision durations is observed during the first 10 minutes, followed by stabilization and then an increase after 15 minutes. Movement times tend to follow an inverse pattern. For monkey B, no significant change in decision or movement durations is observed during the sessions (Figure S1). Overall, this indicates a generally stable effect of the sedation on the animals’ behavior throughout the sessions, which is consistent with the drug’s half-life (1 to 1.2 hours in dogs).

### Relationship between the duration of decisions and movements

We then assessed whether and, if so, how the three monkeys coordinated the duration of their decisions (DD) with the duration of the arm movements (MD) following these decisions to leave the patch they were exploiting.

For monkey G who performed the task under sedation, trial-by-trial analyses indicate a significant correlation between DD and MD for 39 out of 101 sessions (within-session Pearson correlations, p < 0.05). Most of these significant correlations (31/39) are negative, meaning that long decisions were followed by short movements (the “compensation” strategy mentioned in the Introduction). An example session in which compensation occurred in monkey G is shown in Figure 3A. In 8 out of the 101 sessions, a significant positive correlation is observed, indicative of a co-regulation of decisions and movements (i.e., the longest the decision, the longest the movements, and vice versa). Interestingly, correlation coefficients tend to increase across sessions, indicating a progressive switch from a compensation strategy to a co-regulation strategy over time. A linear regression indicates a significant and positive correlation between r-values and session numbers (across sessions Pearson correlation, r=0.33, p=0.0008) (Figure 4, left panel).

**Figure 3-.**
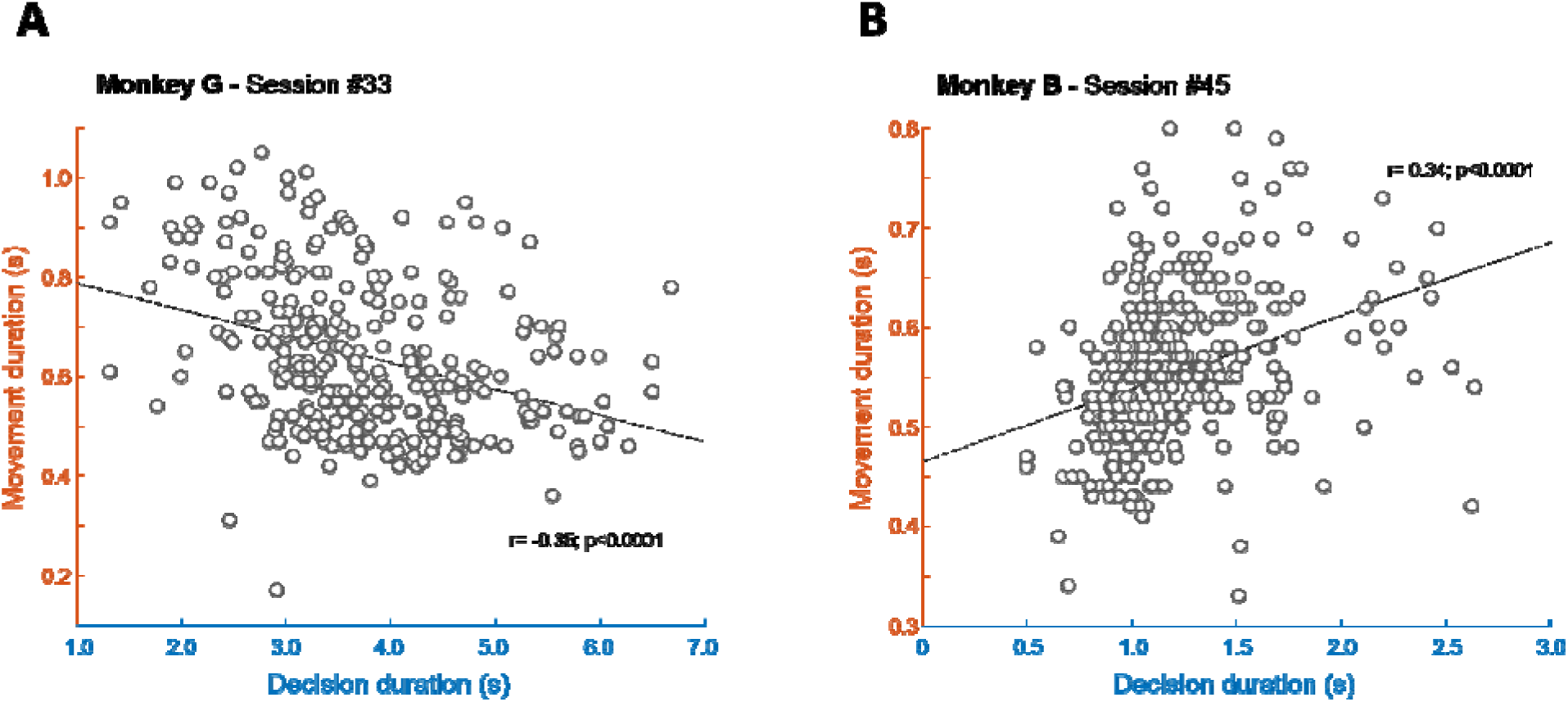
Coordination between decision and movement durations in example sessions. A: Within session, trial-by-trial correlation between decision durations (x-axis) and movement durations (y-axis) for one example session performed by monkey G under sedation condition. Each dot illustrates a trial and the dotted line illustrates the linear regression through the data. B: Same as A for a session performed by monkey B without sedation.

**Figure 4-.**
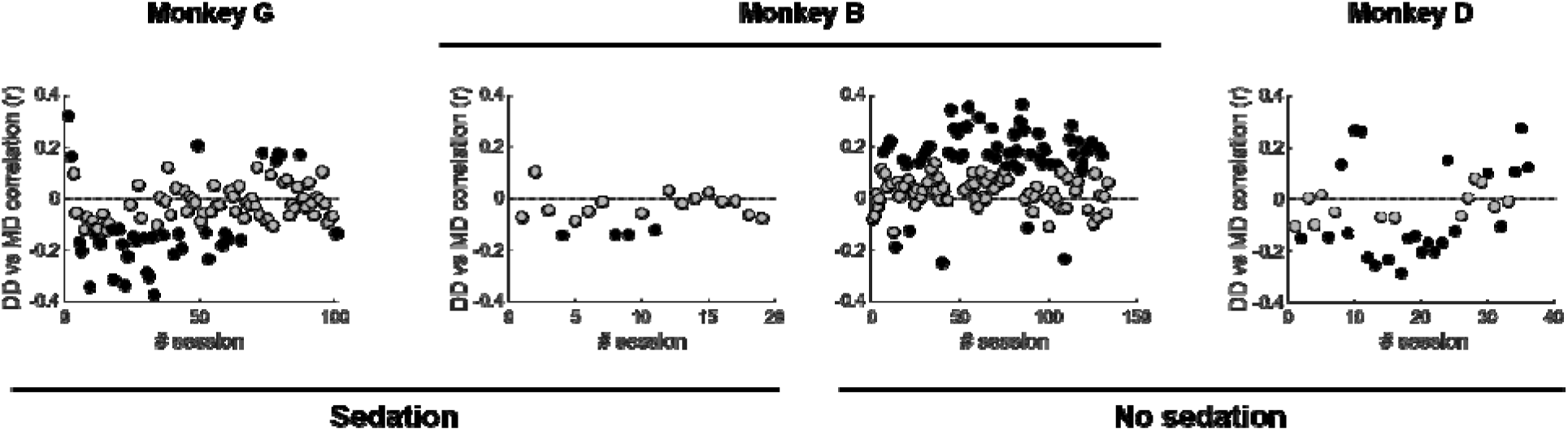
Relation between decision duration and movement duration within sessions. Left panel: Evolution of the trial-by-trial Pearson linear correlation (r) between decision duration (DD) and movement duration (MD) within sessions performed by monkey G under sedation, shown through the time course of these sessions (x-axis). Each dot illustrates the result of a single session. A negative (or positive) r-value indicates that longer decisions are associated with shorter (longer), in terms of duration, movements. Black-filled dots indicate sessions where the Pearson correlation is statistically significant. Second column panel: Same as left for sessions performed by monkey B under sedation. Third column panel: Same as left for session performed by monkey B without sedation. Right panel: Same as left for sessions performed by monkey D without sedation.

When monkey B performed the task without sedation, a significant correlation between DD and MD was observed in 60 out of 133 sessions (within-session Pearson correlations, p < 0.05). Strikingly, almost all of these significant correlations (55/60) are positive, meaning that longer decisions were followed by longer movements, and vice versa (i.e., the “co-regulation” strategy) (Figure 4, third column panel). An example session in which co-regulation occurred in monkey B is shown in Figure 3B. By contrast, when monkey B performed the same task but under sedation, a significant correlation between DD and MD occurred in only 4 out of 19 sessions, and these 4 correlations were negative, indicating compensation between decision and movement durations (Figure 4, second column panel), and were observed primarily during the first half of the sessions performed by the animal.

Finally, for monkey D who performed the task without sedation, a significant correlation between DD and MD is observed in 23 out of 36 sessions (within-session Pearson correlations, p < 0.05). Most of these significant correlations are negative (15/23), indicative of a compensation between decisions and movement durations, but a co-regulation strategy is also observed in 8 sessions.

### Evolution of decision durations, movements durations and reward rates over the course of the sessions

Having observed a change in the coordination between the duration of decisions and movements over the course of the sessions, primarily in monkey G, but also (albeit to a lesser extent) in monkey B under sedation and in monkey D, we sought to determine the extent to which these changes coincided with variations in decision duration, movement duration, or both.

Figure 5 illustrates these changes, showing a large and significant shortening of decision durations (DD) for monkey G over the course of the sessions (across sessions Pearson correlation, r=-0.53, p<0.0001). Average durations were close to 6 seconds during the first 45 sessions but dropped to less than 4 seconds for the final sessions performed by this animal. This trend coincides with a sharp increase in reward rates, at both local and global levels (across sessions Pearson correlations, r=0.58, p<0.0001 and r=0.64, p<0.0001, respectively) (Figure 5, left panels).

**Figure 5-.**
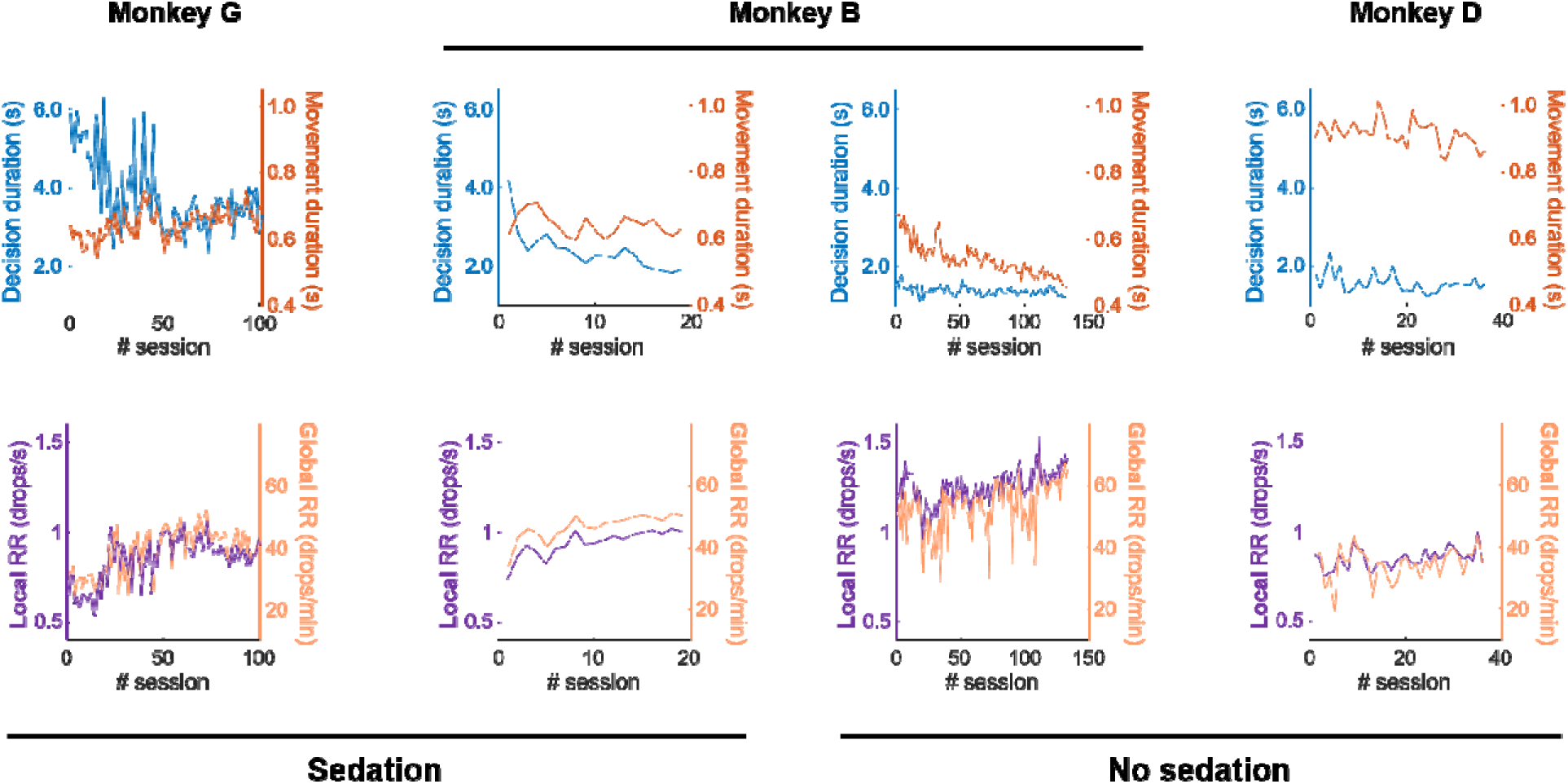
Evolution of decision durations, movement durations and reward rates through the time course of sessions. Left panels: Evolution of the mean decision durations (top, blue), movement durations (top, orange), local (bottom, violet) and global reward rates (bottom, light orange) computed in each session and shown through the time course of these sessions (x-axis) performed by monkey G under sedation. Second column panels: Same as left for session performed by monkey B under sedation. Third column panels: Same as left for session performed by monkey B without sedation. Right panels: Same as left for sessions performed by monkey D without sedation.

A significant decrease in DD is also observed for monkey B under sedation (r=-0.77, p=0.0001), with a reduction from 4 to 2 seconds over the course of the sessions. Here too, an increase in reward rates is observed over the course of the sessions (local reward rate: r=0.83, p<0.0001; global reward rate: r=0.79, p=0.0001) (Figure 5, second column panels).

When monkey B performed the task without sedation, decision times were the shortest and tended to become even shorter across sessions (r=-0.36, p<0.0001). Moreover, a large decrease in movement duration is also noted over the course of the sessions in this condition (r=-0.86, p<0.0001). As a consequence, a gradual increase in reward rates is observed (local reward rate: r=0.56, p<0.0001; global reward rate: r=0.48, p<0.0001) (Figure 5, third column panels).

Finally, for monkey D, a decrease in movement durations is observed over the sessions (r=-0.4, p=0.016), although they remain very long compared to those of the other animals. Decision durations slightly but significantly decreased as well (r=-0.37, p=0.026). Reward rates remained relatively stable throughout the sessions performed by the animal, although they increase significantly when calculated locally (local reward rate: r=0.41, p=0.01; global reward rate: r=0.24, p=0.15) (Figure 5, right panels).

## Discussion

The present study provides causal evidence that the coordination between decision duration and movement duration is not a fixed property of behavior but a flexible process that adapts to changes in the animal’s internal state. By pharmacologically slowing decision formation while leaving the behavioral task unchanged, we induced a transient reorganization of the temporal relationship between deliberation and movement execution. Initially, prolonged decisions were predominantly associated with shorter movements, consistent with a compensatory strategy limiting the additional temporal cost imposed by slower deliberation. As animals progressively adapted to the perturbation, decision durations decreased, reward rates increased, and compensatory coordination became less prevalent, giving way to patterns of decision-movement uncoupling (monkey B) or co-regulation (monkey G). Together, these findings suggest that the coordination between decision-making and movement execution contributes to preserving behavioral efficiency when internal perturbations alter the temporal organization of behavior.

Our previous work, in both monkeys and healthy humans, together with studies from other groups, has shown that decision duration and movement vigor influence one another under a wide range of behavioral contexts (Thura et al., 2014; Reynaud et al., 2020; Carsten et al., 2023; Kaduk et al., 2023; Kita et al., 2023; Saleri Lunazzi et al., 2023; Fievez et al., 2024; Morvan et al., 2024; Saleri and Thura, 2024; Leroy et al., 2025; Saleri et al., 2026). However, these demonstrations were based almost exclusively on manipulations of external variables, including changes in movement requirements, decision difficulty, or reward contingencies. Such manipulations leave open the possibility that coordinated adjustments emerge simply because individuals adapt to altered task constraints rather than because they intrinsically and actively regulate the temporal organization of decisions and actions. The present study addresses this limitation by perturbing the animal’s internal state while maintaining the external environment unchanged. The observation that a mild pharmacological slowing of decision formation was accompanied by systematic changes in the relationship between decision duration and movement duration therefore provides stronger support for the existence of an integrated control process coordinating these two components of behavior (Yoon et al., 2018; Sukumar et al., 2024).

Importantly, the compensatory adjustments observed under sedation cannot be readily interpreted as a trivial motor consequence of the pharmacological treatment. In monkey B, sedation resulted in an 18% increase in MD, while DD increased by 41%. If sedation simply slowed the motor system (in addition to decision-making), one would expect longer decisions to be accompanied by longer movements, thereby reinforcing the temporal consequences of the perturbation. Instead, the opposite tendency predominated during the early stages of adaptation: prolonged deliberation was preferentially followed by shorter movements (Figure 3A). Such compensation is difficult to reconcile with a passive effect of the drug on both decision-making and movement execution but is consistent with an active behavioral adjustment aimed at limiting the overall temporal cost of each foraging trial. Although the present data do not identify the neural mechanisms responsible for this adjustment, they provide causal evidence that modifying decision duration is sufficient to reshape the coordination between decision-making and movement execution.

These observations further strengthen normative accounts proposing that decisions and movements should not be optimized independently but rather as successive components of a common behavioral sequence (Yoon et al., 2018; Sukumar et al., 2024). In natural behavior, both deliberation and movement contribute to the duration separating two successive rewards (Cisek, 2007; Cisek and Kalaska, 2010). Consequently, a prolongation of either component inevitably reduces the rate at which rewards can be acquired unless compensated by adjustments in the other. The compensatory coordination observed here is fully consistent with this theoretical framework (Bogacz et al., 2010; Balci et al., 2011). Rather than attempting to immediately restore normal decision times, the animals initially appeared to compensate for part of the additional time cost associated with decision-making by modifying the execution of subsequent movements. Such adjustments do not completely eliminate the consequences of slower decisions (monkey B’s reward rates are lower with sedation compared to without sedation) but reduce their impact on overall behavioral efficiency.

The progressive increase in both local and global reward rates further supports this interpretation. As animals adapted to the sedative perturbation (decision durations of both monkey G and monkey B progressively decreased across sessions under sedation), reward rates increased toward values observed under control condition (for monkey B). Although reward rate maximization cannot be demonstrated directly from the present data, the observed adjustments are consistent with a control policy aimed at preserving the efficiency of reward acquisition despite a transient degradation of deliberation duration. From this perspective, the coordination between decisions and movements may represent one component of a broader adaptive strategy through which the nervous system regulates the temporal organization of behavior in response to changing internal conditions.

One of the most interesting observations of the present study is that compensatory coordination was not stable across the course of adaptation. Instead, the negative relationship between the duration of decisions and movements progressively weakened as animals recovered from the pharmacological perturbation across sessions, while patterns of positive decision-movement co-regulation became increasingly prevalent (at least for monkey G). This observation suggests that compensatory coordination is not a constitutive feature of behavior but rather a transient response that emerges when the temporal organization of behavior is challenged.

At first glance, the replacement of compensation by co-regulation may appear inconsistent with the hypothesis that decision and movement are jointly regulated to maximize reward rate. However, this apparent contradiction disappears if one considers that the objective of the control policy may be to maintain efficient behavioral performance under changing internal conditions. During the early stages of sedation, when decisions were substantially prolonged, reducing movement duration represented one possible means of limiting the temporal cost imposed by slower deliberation. As animals progressively adapted and decision durations returned toward baseline values, this compensatory adjustment became less necessary. Under these “natural” conditions, the temporal relationship between decisions and movements may once again become dominated by processes that influence both components in the same direction. It is interesting to note that this mode of regulation seems to operate even when not required, when subjects cannot control their speed/accuracy trade-off (Kita et al., 2023; Fievez et al., 2024). This suggests that the co-regulation of decisions and movements constitutes a default mode of coordination, possibly resulting from neural circuits shaped by evolution and designed to respond effectively to most real-life urgent situations requiring both fast decisions and vigorous actions (Carsten et al., 2023; Fievez et al., 2024; Saleri and Thura, 2024; Thura et al., 2025).

Several mechanisms could account for this co-regulation. Previous work has suggested that decision urgency, motivational state, or the subjective valuation of time may jointly influence deliberation and movement vigor (Thura et al., 2014, 2025; Manohar et al., 2015; Shadmehr et al., 2016; Thura and Cisek, 2016; Shadmehr and Ahmed, 2020; Thura, 2020; Korbisch et al., 2022; Kaduk et al., 2023). Fluctuations in these internal variables would naturally produce positive covariations between decision and movement durations without requiring an explicit and costly compensatory process. Rather than reflecting two qualitatively distinct coordination strategies, compensation and co-regulation may therefore correspond to different behavioral expressions of a common adaptive controller operating under different constraints.

This interpretation is also consistent with the marked improvement in reward rate observed throughout adaptation. Importantly, behavioral efficiency increased primarily because decision durations progressively shortened rather than because compensatory coordination itself became stronger. Compensation therefore appears to provide a temporary solution that limits the impact of an internal perturbation while the behavioral system gradually recovers a more efficient operating regime. In this view, coordinated adjustments between decision and movement are not an end in themselves but one component of a broader adaptive process through which behavior remains robust despite transient changes in internal state or in external constraints.

It is also very interesting to note that these adaptive mechanisms are not limited to responding to perturbations in decision-making. Indeed, in the case of monkey D who produced extremely long movements for reasons of his own (the monkey was not sedated during the experimental sessions), the prevailing mode of coordination was compensation. Interestingly, these movement durations tended to decrease over the course of the sessions, and the coordination then appeared to reflect, to a greater extent, a co-regulation between the two processes.

Several limitations should nevertheless be considered when interpreting the present findings. First, only one animal was tested both with and without sedation, whereas monkey G was always recorded under the sedative condition. Although the convergence between animals strengthens the general interpretation, additional within-subject manipulations would provide a more direct assessment of the causal effects of pharmacological slowing. Second, the sedative doses differed slightly between monkeys, reflecting experimental constraints rather than a systematic manipulation of drug concentration. Finally, the progressive shortening of decision durations likely reflects a combination of behavioral adaptation, continued task experience, and possibly changes in sensitivity to repeated pharmacological treatment. Disentangling these factors will require future experiments specifically designed to manipulate internal state independently of learning.

Despite these limitations, the present results extend previous demonstrations of decision-movement interactions by showing that an experimentally induced perturbation of the animal’s internal state is sufficient to reorganize the temporal coordination between deliberation and movement execution. More broadly, they support the view that decisions and movements should not be regarded as independent stages of behavior, but as complementary components of a unified adaptive control process that flexibly regulates the temporal organization of behavior in response to internal perturbations (Thura et al., 2022). Understanding how the nervous system implements this integrated regulation may therefore provide important insights into the principles governing adaptive behavior across changing internal and external environments.

## Acknowledgements/Funding

The authors thank Sonia Alouche for her administrative assistance, Frédéric Volland for his contribution during the technical preparation and maintenance of this experiment, Michel Filiptchenko and Manon Dirheimer for their daily care of the monkeys in our animal facility. They also thank Luc Renaud who performed the surgical procedures mentioned in this study. ChatGPT (chatgpt.com) and Google Translate (translate.google.com) were used to correct the English, without altering the content. This work is supported by an ANR grant (ANR-22-CE37-0010-02/ BasalCost) to DT.

