## Supplemental Figure 1 for "Internal perturbation reveals the flexible and adaptive nature of the coordination between decisions and movements"

*Supplemental information*

### Supplemental figure

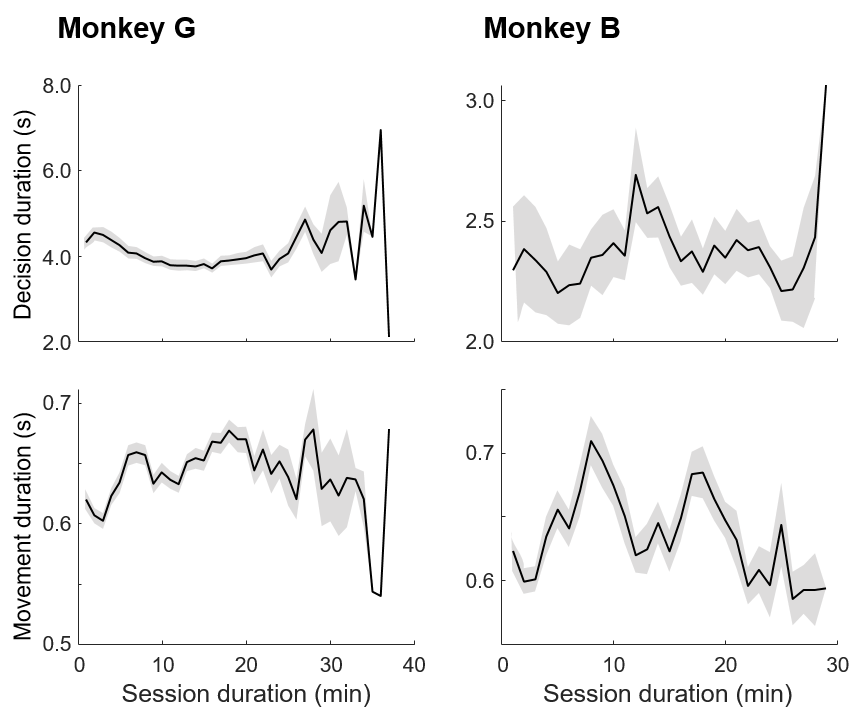

**Figure S1: Evolution of decision durations (top) and movement durations (bottom) of monkey G (left) and monkey B (right) within sessions performed under sedation.** The black curves represent the mean values calculated across all sessions, with standard errors indicated by the shaded areas. The values are calculated for each minute of the task performed by the animals, with time corresponding to the total duration of the trials included in the analyses presented in the main paper.
